# Amblyopic deficits in monocular processing and binocular interactions revealed by submillimeter 7T fMRI and EEG frequency tagging

**DOI:** 10.1101/2024.04.25.591076

**Authors:** Yue Wang, Chencan Qian, Yige Gao, Yulian Zhou, Xiaotong Zhang, Wen Wen, Peng Zhang

## Abstract

Disruption of retinal input early in life can lead to amblyopia, a condition characterized by reduced visual acuity despite corrected optics. Although extensive losses of neural activity have been found in the early visual cortex, it remains unclear whether they reflect deficits in feedforward or feedback processing, or abnormal binocular interactions. Combining submillimeter 7T fMRI and EEG frequency tagging, our study revealed the precise neural deficits in monocular processing and binocular interactions in human adults with unilateral amblyopia. Cortical depth-dependent fMRI revealed monocular response deficits in cortical layers of the primary visual cortex (V1) receiving thalamic input, which carried over to the downstream areas (V2-V4) in feedforward processing. Binocular stimulation produced a greater signal loss in the superficial layers of V1, consistent with suppression from the fellow eye by lateral inhibition. EEG data further demonstrate reduced suppression from the amblyopic eye, weakened binocular integration, and delayed monocular and binocular processing. Our results support attenuated and delayed monocular processing in V1 layers receiving thalamic input in human amblyopia, followed by imbalanced binocular suppression and weakened binocular integration in the superficial layers, which further reduced signal strength and processing speed. These precise neural deficits can help developing more targeted and effective treatments for the vision disorder.

**Significance:** Cortical depth-dependent 7T fMRI and EEG frequency tagging revealed attenuated and delayed neural activity in monocular processing and binocular interactions in human amblyopia. Neural deficits in monocular processing arise from the input layers of V1, followed by imbalanced binocular suppression and weakened binocular integration in the superficial layers. The precise neural deficits at high spatiotemporal resolution can help developing more targeted and effective treatment for amblyopia.

**Graphical abstract:** 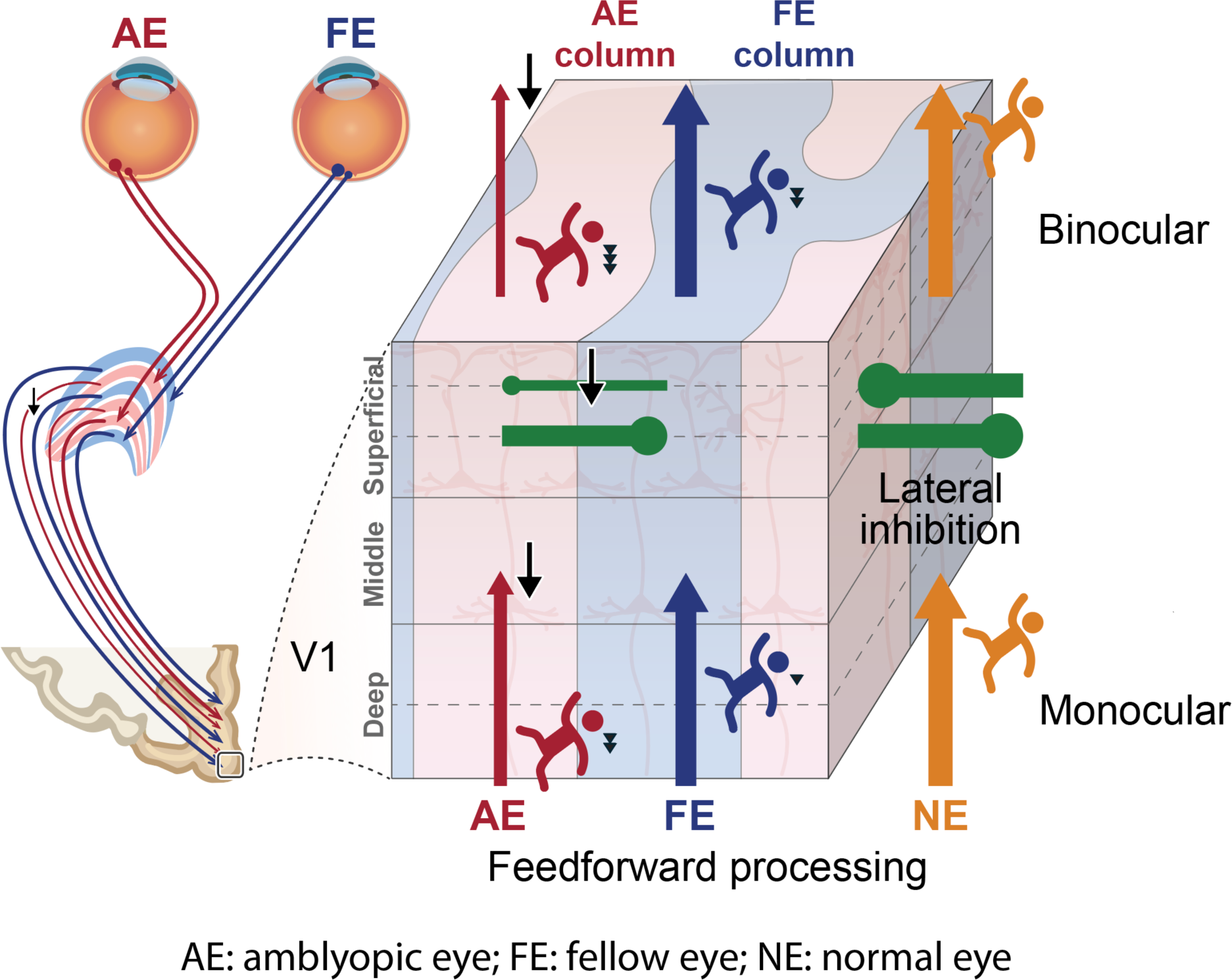

This figure illustrates the precise neural deficits of amblyopia in monocular processing and binocular interactions based on the main findings of the current study. Attenuated and delayed neural activity arises from V1 cortical layers receiving thalamic input in monocular processing, followed by imbalanced binocular suppression and weakened binocular integration in the superficial layers, further reducing visual signal strength and processing speed.

## Introduction

Amblyopia, commonly known as “lazy eye”, is a neurodevelopmental disorder characterized by visual acuity loss due to early-life disruptions of retinal input, such as anisometropia (unequal refractive error) or strabismus (misalignment of eyes). Understanding the precise neural deficits of amblyopia is crucial for developing more targeted and effective therapeutic interventions. Behavioral and neuroimaging studies have found monocular deficits in both low- and high-level visual functions (Levi, 2020). However, it remains unclear whether they reflect abnormalities in feedforward or feedback processing. For example, high-level deficits could be simply inherited from low-level areas, and may also affect the activity of these regions through feedback modulation. In addition to monocular deficits, clinical and psychophysical studies also suggest abnormalities in binocular suppressions (Hess et al., 2014; Zhou et al., 2018), consistent with electrophysiological evidence in animal models (Bi et al., 2011; Shooner et al., 2017). However, functional abnormalities in binocular interactions remain highly controversial in neuroimaging studies of human amblyopia (Baker et al., 2015; Chadnova et al., 2017; Farivar et al., 2011; Lygo et al., 2021).

The primary visual cortex (V1) consists of six layers of neurons that play distinct roles in feedforward, feedback and intracortical processing (Felleman & Van Essen, 1991). As illustrated by the laminar circuitry in V1 (Fig. 1a), input from the lateral geniculate nucleus (LGN) of the thalamus terminates mainly in the middle layer (4c) and with a smaller portion in the deep layer (6). The deep (5/6) and superficial layers (1/2/3) receive feedback projections from the higher cortex. Horizontal or lateral inhibitions between ocular dominance columns (ODCs) are most prominent in the superficial layers (2/3) as demonstrated by electrophysiological recordings and intrinsic axonal connections (Cox et al., 2019; Gilbert & Wiesel, 1983; Sengpiel et al., 1995). Recent development of high-resolution fMRI at ultra-high magnetic fields (7 Tesla and above) allows to non-invasively measure cortical depth-dependent activity in humans (Fracasso et al., 2018; Huber et al., 2017; Kok et al., 2016; Liu et al., 2021), making it practically possible to study mesoscale functional abnormalities in cortical microcircuits.

**Figure 1.**
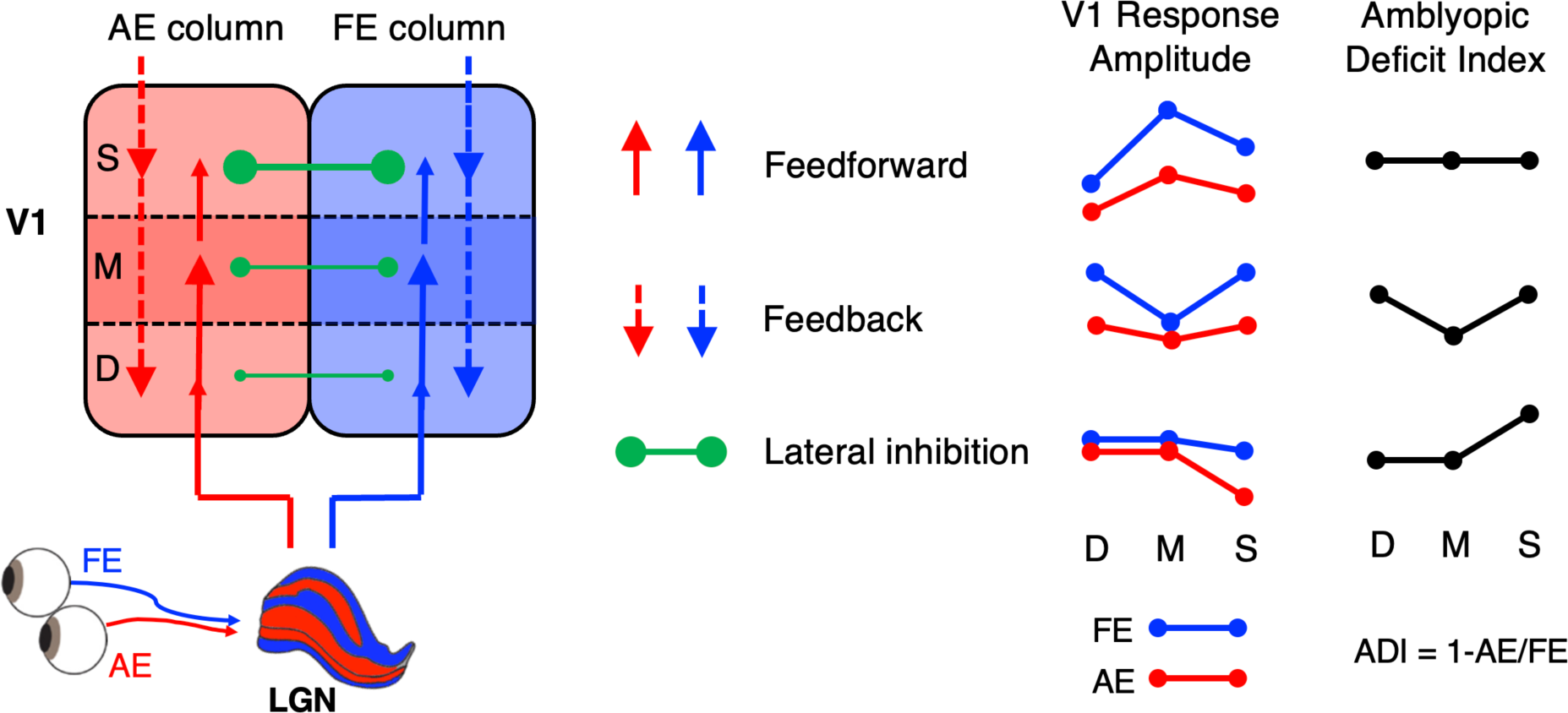
**(a)** Neural circuits in the geniculostriate pathway that may be affected by amblyopia. Solid and dashed arrows denote feedforward and feedback connections, respectively. Green dots connected by a solid line indicate lateral inhibitions between adjacent ocular dominance columns. **(b)** Layer-specific response and amblyopic deficit in V1. Red and blue lines indicate AE and FE responses, respectively. The amblyopic deficit index (ADI) was defined as the proportion of response loss in AE compared to FE. Abbreviations: S: superficial layers, M: middle layers, D: deep layers, AE: amblyopic eye. FE: fellow eye.

In experiment 1 (Exp. 1), to investigate the cortical microcircuits underlying the neural deficits of amblyopia, we used submillimeter resolution (0.8 mm isotropic) fMRI at 7 Tesla to measure layer-dependent responses in V1 of 10 human adults with unilateral amblyopia (see Table S1 for clinical characteristics). As in previous electrophysiological studies in animal models (Kiorpes et al., 1998; Shooner et al., 2017), we defined an amblyopic deficit index (ADI) by the response difference between the amblyopic eye (AE) and the fellow eye (FE) divided by the FE response (Fig. 1b). If the AE activity loss arises from the thalamic input and propagates along the feedforward pathway, ADI would be comparable across cortical depths. If higher cortical deficits affect V1 processing through feedback modulations, ADI would be evident in the superficial and deep layers. Finally, in the binocular stimulus condition, stronger suppression from FE to AE than vice versa would produce a larger response deficit in the superficial layers of V1 by lateral inhibition.

In experiment 2 (Exp. 2), to further characterize the amblyopic abnormalities in binocular interactions and visual processing speed, we used an EEG frequency-tagging method to measure steady-state visually evoked potentials (SSVEPs) to stimuli presented to the two eyes at slightly different temporal frequencies (Norcia et al., 2015; P. Zhang et al., 2011). A total of 37 adults with unilateral amblyopia (see Table S1 for clinical characteristics) and 15 healthy controls participated in the EEG experiment.

## Results

In the 7T fMRI experiment, full contrast checkerboard flickers were presented to the amblyopic eye (AE), fellow eye (FE) or both eyes (binocular) in separate blocks (Fig. 2a). Subjects viewed the stimuli through a pair of MRI-compatible goggles (Fig. 2b). A 32-channel receive 4-channel transmit surface coil was used to acquire high-quality MRI data from the occipital and temporal lobes (Sengupta et al., 2016). A bite bar was used to reduce head motion. Fig. 2c shows the segmentations of cortical gray and white matter in a representative subject. Fig. 2d shows the visually evoked activation maps to AE, FE and binocular stimuli, and AE-FE response for the same subject.

**Figure 2.**
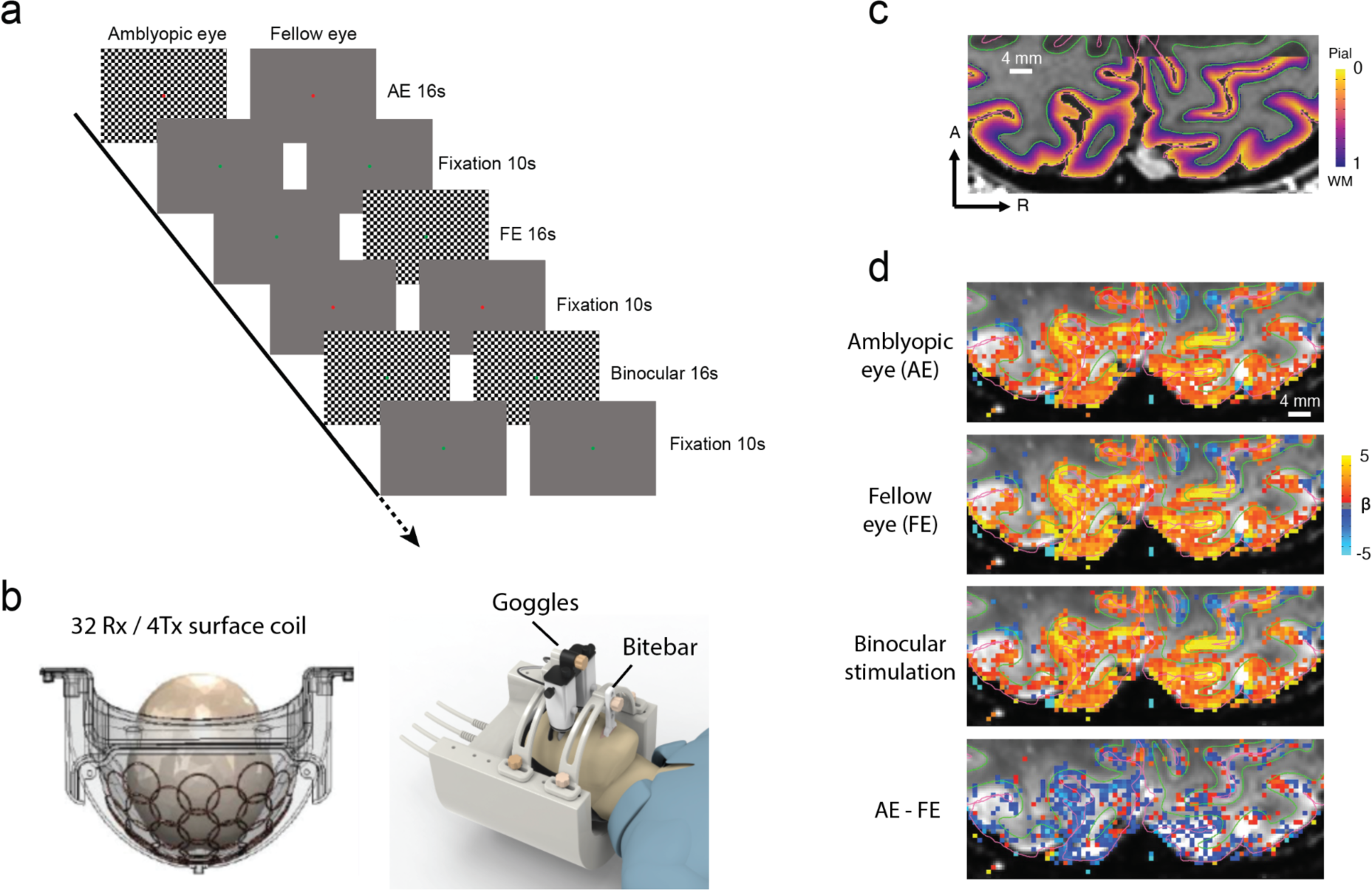
**(a)** Stimuli and procedure of the fMRI experiment. **(b)** The open-face surface coil, goggles, and bite bar. **(c)** Cortical depth map and segmentation in the occipital lobe of a representative subject (P10). Green and purple lines denote the white matter and gray matter surfaces. The underlay shows the structural image (T1-MP2RAGE). **(d)** Occipital activations to visual stimuli in the same subject. The underlay shows the mean functional image (GE-EPI). Maps were thresholded at p < 0.5, uncorrected.

### Amblyopic response deficits in different cortical depths of V1 and in higher visual areas

BOLD responses in V1 showed a strong bias toward the superficial depth in both monocular and binocular stimulus conditions (Fig. 3a), consistent with influence from intracortical draining veins and large pial veins (Havlicek & Uludağ, 2020; Huber et al., 2019). However, the amblyopic deficit index (ADI) showed distinct laminar profiles in the monocular and binocular stimulation conditions (Fig. 3b), supported by a highly significant interaction between ocularity (monocular/binocular) and layers (superficial/middle/deep) in a two-way repeated measures (rm) ANOVA (F(2,18) =19.068, *p* < 0.001, η^2^_p_= 0.680). Monocular ADIs were comparable across cortical depths (F(2,18) = 1.519, p = 0.246, BF_01_ = 1.885), while binocular ADI was strongest at the superficial depth (F(2,18) = 14.224, p < 0.001, η^2^_p_ = 0.612; S vs. M: t(9) = 4.250, p = 0.002, Cohen’s d = 1.002; S vs. D: t(9) = 4.275, p = 0.002, Cohen’s d = 1.361). To avoid double dipping, monocular responses were extracted from visually responsive voxels within an anatomically defined V1 ROI, while the binocular responses were extracted from voxels showing strong ocular bias in the monocular stimulation conditions (see Methods; see Fig. S2 for monocular and binocular results from the same voxels using a split-half approach).

**Figure 3.**
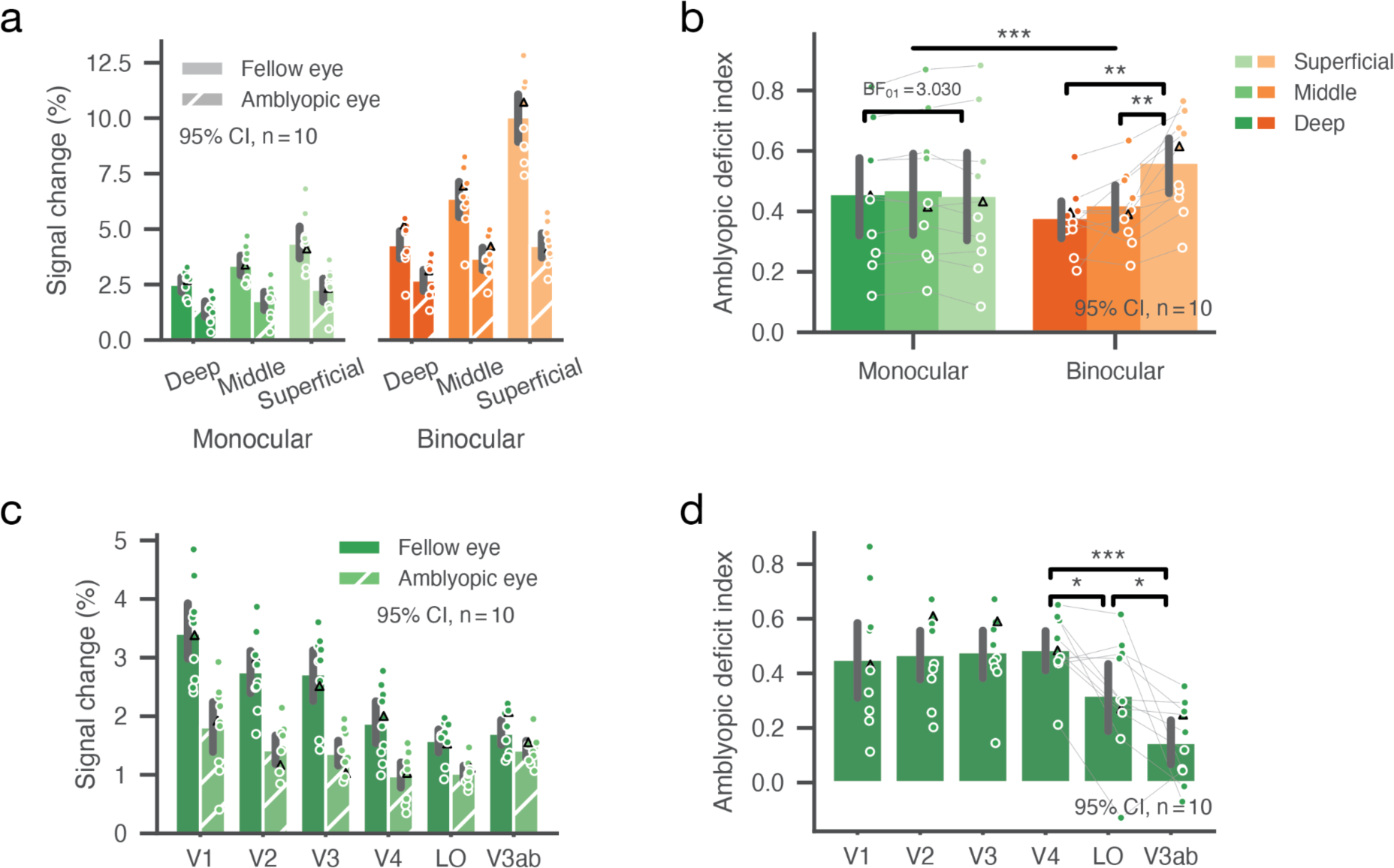
**(a)** Cortical depth-dependent fMRI response in V1. **(b)** Amblyopic deficit indices in the monocular and binocular stimulus conditions across cortical depths in V1. **(c)** BOLD responses to monocular stimuli in V1 and higher cortical regions. **(d)** Monocular ADIs in the visual cortical areas. In all panels, each dot represents datum from one subject (circle for anisometropia, triangle for strabismus). *, **, and *** denote p < 0.05, 0.01 and 0.001, respectively. Error bars represent 95% confidence intervals.

Monocular response deficit was also found in higher cortical regions beyond V1 (Fig. 3c). Monocular ADIs were comparable in the early visual areas from V1 to V3 (Fig. 3d; F(2,18) = 0.131, p = 0.878, BF_01_ = 4.333), but there was a significant ventral-to-dorsal gradient of alleviated deficits in the middle visual areas V4, LO, and V3ab (F(2,18) = 14.710, p < 0.001, η^2^_p_= 0.620; V4 vs. LO: t(9) = 3.065, p = 0.013, Cohen’s d = 0.969; V4 vs. V3ab: t(9) = 5.842, p <0.001, Cohen’s d = 2.559; LO vs. V3ab: t(9) = 2.349, p = 0.043, Cohen’s d =0.968). These findings support that the amblyopic deficit in monocular processing arises from the thalamic input and carries over to the downstream areas along the feedforward pathway (Fig. 1), and the dorsal pathway is more resilient to the degraded inputs, especially in high spatial frequency components. In the binocular viewing condition, stronger suppression from the fellow eye leads to an additional response loss in the superficial layer of V1 by lateral inhibition.

Due to the partial volume effect of fMRI, voxels biased to one eye were influenced by signals from the opposing eye, even at the submillimeter resolution. This can lead to an underestimation of binocular ADIs in Fig. 3b, suggesting that the binocular ADI would be even larger in the superficial depth of V1 (Fig. 4c). However, the partial volume effect poses a challenge to measure the effect of binocular suppression, since the binocular response could be stronger than the monocular response even in the ocular biased voxels. To further investigate the effect of binocular suppression and its relationship with the amblyopic deficits, we used an SSVEP frequency-tagging method in Exp. 2 to measure the two eyes’ responses to naturalistic stimuli presented at slightly different temporal frequencies. Visual stimuli were low- and high-pass filtered (cutoff frequency at 2 c.p.d.) to selectively evoke neural activity biased to the magnocellular (M) and parvocellular (P) pathways, respectively.

**Figure 4.**
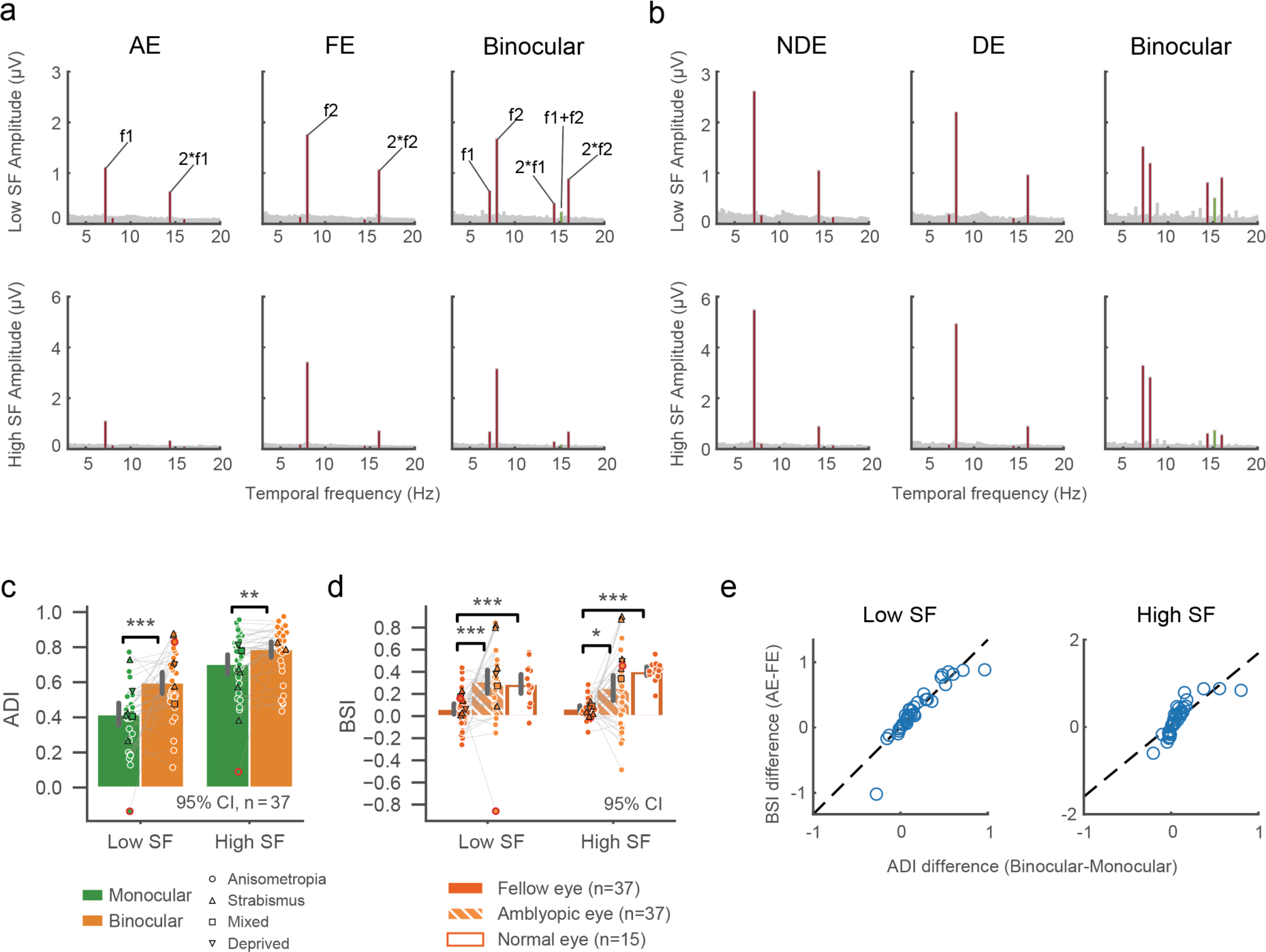
**(a)** Amplitude spectrums of SSVEPs in monocular and binocular conditions averaged across all amblyopic subjects. The stimulus frequencies in AE and FE were f1 = 7.2 Hz, and f2 = 8 Hz, respectively. Fig. S3 shows results from the opposite setup (f2 in AE and f1 in FE). Green lines represent the intermodulation frequency at f1+f2. Upper and lower rows show the results for low and high spatial frequency (SF) stimuli, respectively. **(b)** SSVEP amplitude spectrum in normal controls (f1 in non-dominant eye (NDE) and f2 in dominant eye (DE); See Fig. S4 for f2 in NDE and f1 in DE). **(c)** Amblyopic deficit index (ADI) for both monocular (green) and binocular (orange) stimulus conditions. Different symbols represent individuals with different types of amblyopia: anisometropia (circle), strabismus (triangle up), mixed (square), deprived (triangle down). Symbols with red outline are detected as outlier based on within-group multivariate Mahalanobis distance (see Methods) and excluded from reported statistics. Error bars represent 95% confidence intervals. *p < 0.05, **p < 0.01, ***p < 0.001. (**d**) Binocular suppression index (BSI) for the FE (solid), AE (dashed), and normal eye (NE, unfilled), otherwise same convention as in (c). **(e)** Correlations between the difference in ADI (binocular-monocular) and in BSI (AE-FE). Open circles represent data from individual subjects.

### Reduced suppression from AE correlates with binocular response deficits

As shown in Fig. 4a/b, the amplitude spectrum of SSVEPs in both amblyopic and control groups showed high signal-to-noise ratio at the stimulus frequencies. Binocular ADIs were significantly larger than monocular ADIs (Fig. 5c), demonstrated by a significant main effect of ocularity (F(1,35) = 18.898, p < 0.001, η^2^_p_= 0.351) in a two-way rm ANOVA. This effect was significant in both low (t(35) = −4.336, p < 0.001, Cohen’s d = −0.895) and high (t(35) = - 3.027, p = 0.005, Cohen’s d = −0.431) spatial frequency (SF) conditions, but more robust in low compared to high SF (F(1,35) = 9.491, p = 0.004, η^2^_p_= 0.213). We also computed a binocular suppression index (BSI), defined as the amplitude difference between monocular and binocular conditions divided by the monocular amplitude (1-binocular/monocular), to indicate the degree of suppression by the opposing eye (Fig. 4c). For patients with amblyopia, BSI was significantly larger for AE compared to FE (F(1,35) = 22.213, p < 0.001, η^2^_p_= 0.388), in both low (Wilcoxon signed-rank test W = 44, p < 0.001, rank-biserial correlation (RBC) = −0.868) and high (W = 162, p = 0.019, RBC = −0.514) SF conditions. In healthy controls, BSIs showed no significant difference between the dominant (DE) and non-dominant (NDE) eyes, thus were averaged as the normal eye (NE) group. In the low SF condition, BSI for NE was significantly larger than FE (Wilcoxon rank-sum test U = 84, p < 0.001, RBC = −0.689) but close to AE (U = 310, p = 0.414, RBC = 0.148). Similarly in the high SF condition, BSI for NE was significantly larger than FE (U = 3, p < 0.001, RBC = - 0.989) but not than AE (U = 170, p = 0.079 after Holm correction, RBC = −0.370).

**Figure 5.**
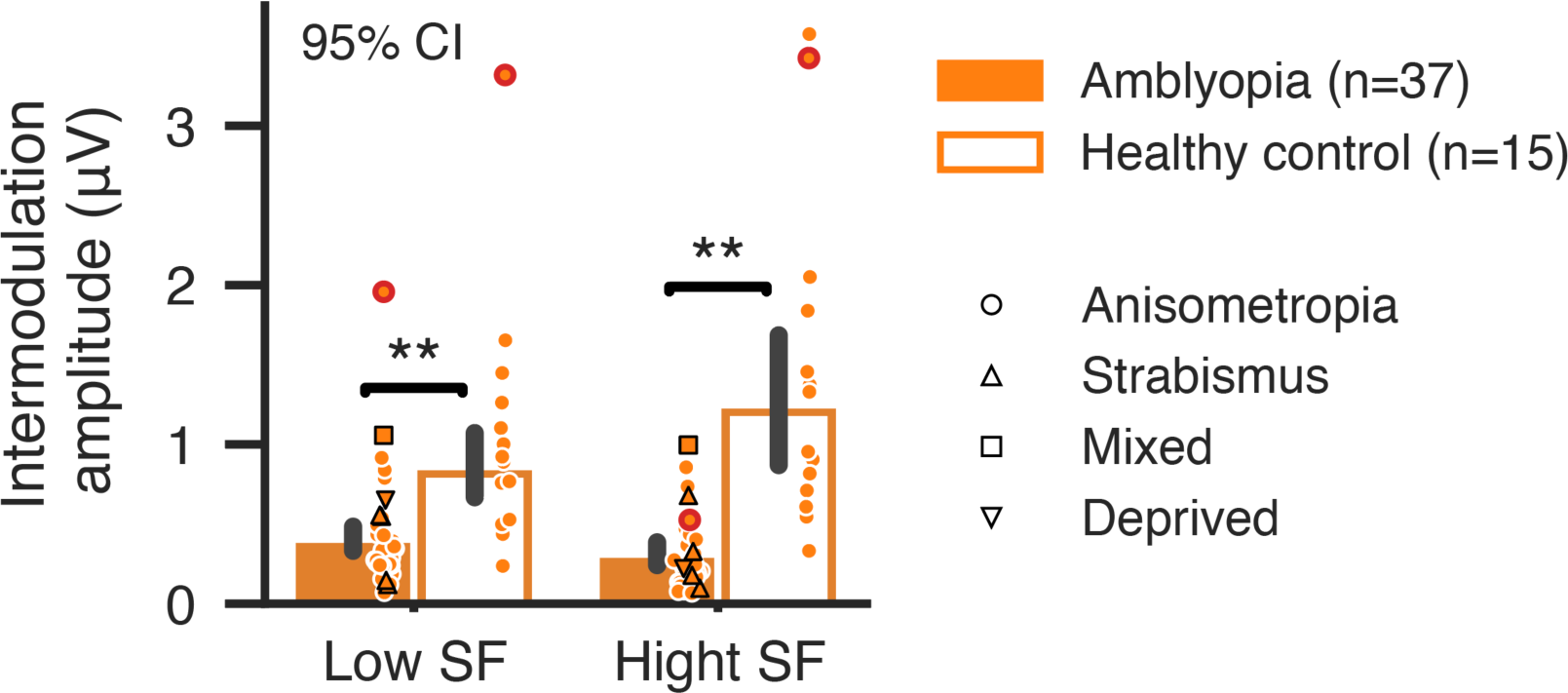
SSVEP amplitudes at the intermodulation frequency (f1+f2). Same conventions as in Fig. 4c/d.

These findings provide strong evidence for asymmetric binocular suppressions between AE and FE, i.e. stronger suppression from FE to AE than vice versa. This was mainly due to reduced suppression by AE compared to binocular suppressions in the control group. To further investigate whether the asymmetric binocular suppression is related to the additional deficits in the binocular condition, we calculated Pearson’s correlation between the difference in ADI (binocular-monocular) and in BSI (AE-FE). Results showed a highly significant correlation in both low (r = 0.895, p < 0.001) and high (r = 0.743, p < 0.001) SF conditions.

### Reduced intermodulation amplitude indicates weakened binocular integration

To further investigate functional abnormalities in binocular integration, we examined the difference in the amplitude of intermodulation (IM) frequency (f1+f2) between amblyopes and healthy controls (Fig. 5). IM amplitudes were significantly reduced in the amblyopic group (F(1,48) = 44.601, p < 0.001, η^2^_p_= 0.482), and in both high and low SF conditions (high SF, t(13.784) = −4.159, p = 0.002, Cohen’s d = −1.973; low SF, t(16.384) = −3.946, p = 0.002, Cohen’s d = −1.572). Moreover, there was a significant SF × group interaction (F(1,48) = 12.901, p < 0.001, η^2^_p_= 0.212) in IM amplitude, suggesting larger deficit in binocular integration in the high SF condition.

### Delayed neural activity to monocular and binocular stimuli in the early visual cortex

To investigate the amblyopic deficits in visual processing speed, we analyzed the phase of SSVEPs in AE, FE and NE conditions using a least-squares filter method (Tang & Norcia, 1995). The group averaged SSVEP timecourses in Fig. 6a show clear phase delays between AE and FE, and between FE and NE. In addition, neural activity to high SF stimuli was also clearly delayed compared to the low SF activity.

**Figure 6.**
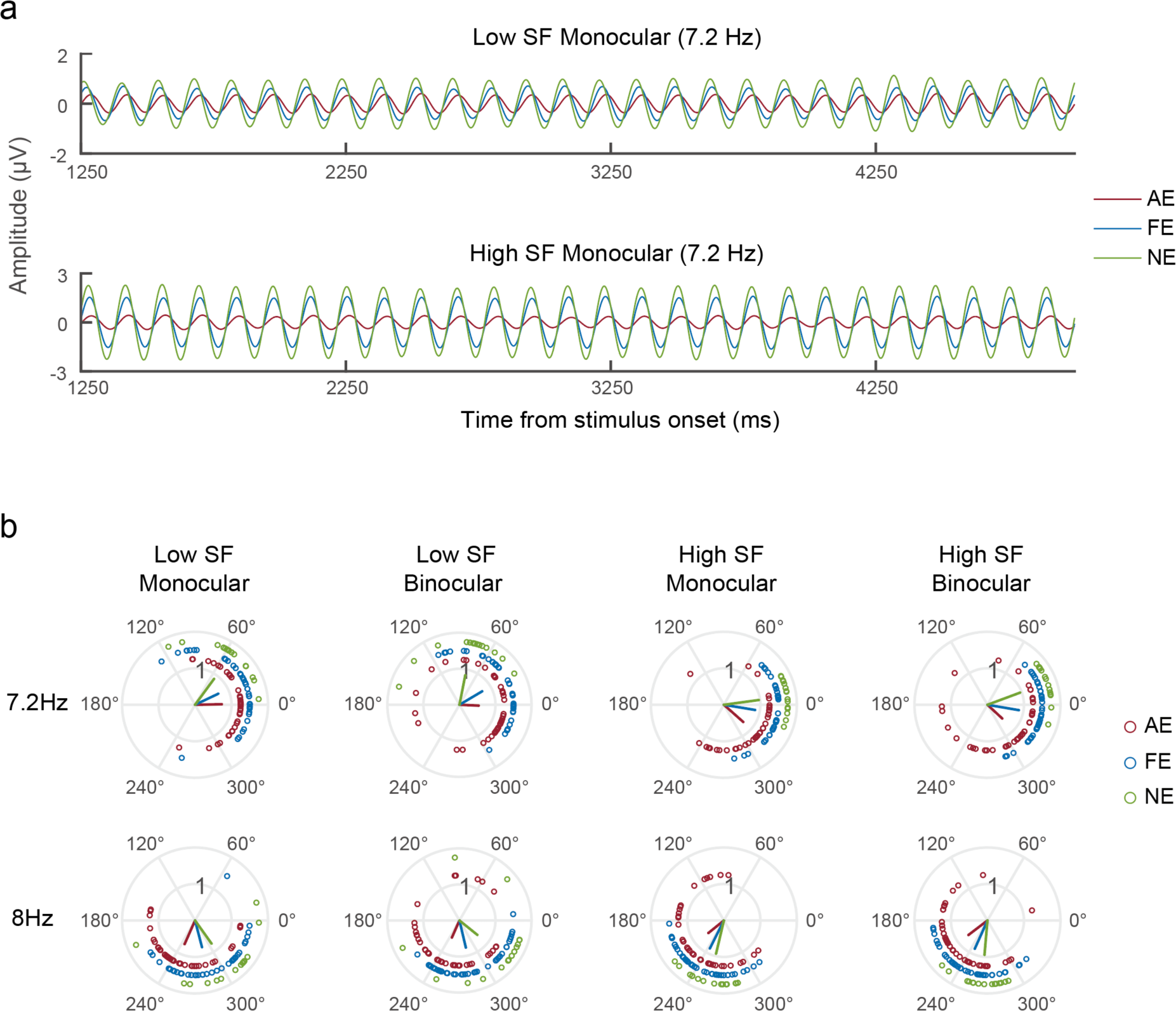
**(a)** Group averaged SSVEPs. The monocular stimulus conditions at 7.2 Hz were shown here as examples (from 1250 to 5000 ms after stimulus onset). **(b)** Polar plots of SSVEP phases. Red, blue and green lines denote group averaged phases for AE, FE and NE responses. Circles represent individual data. DE and NDE phases were averaged for the NE group.

The polar plot in Fig. 6b illustrates the phase of SSVEPs in all stimulus conditions. Circular statistics revealed a significant phase delay between AE and FE (circ_mtest, p < 0.001, mean delay = 12.81 ms), and between AE and NE (Watson-Williams test, p < 0.001, mean delay = 21.69 ms) (Berens, 2009). FE responses were also significantly delayed relative to NE (Watson-Williams test, p = 0.003, mean delay = 8.88 ms). Neural responses to binocular stimuli were also significantly faster compared to the monocular stimulus conditions in normal controls (circ_mtest, p < 0.001; mean delay = −6.24 ms), but not in amblyopic subjects (AE: p = 0.237, mean delay = −0.98 ms,; FE: p = 0.721, mean delay = −0.089 ms). This aligns with our finding of impaired binocular integration indexed by reduced intermodulation amplitude. Finally, high SF responses were significantly delayed compared to the low SF conditions (circ_mtest, FE: p < 0.001, mean delay = 14.27 ms; NE: p <0.001, mean delay = 20.87 ms; AE: p = 0.022, mean delay = 9.27 ms), consistent with slower processing speed of the parvocellular pathway.

## Discussion

Using submillimeter 7T fMRI and EEG frequency tagging methods, we investigated the precise neural deficits in monocular processing and binocular interactions in human adults with unilateral amblyopia. Monocular response deficits in V1 were comparable across cortical depths and similar to those in the extrastriate visual areas (V2-V4), suggesting amblyopic deficit in cortical layers receiving thalamic input that are inherited by downstream visual areas in feedforward processing. In the binocular stimulus condition, a stronger response deficit was found in the superficial layers of V1, consistent with suppression by the fellow eye through lateral inhibition. SSVEPs further demonstrate reduced suppression from the amblyopic eye, weakened binocular integration, as well as delayed monocular and binocular processing. These findings provide robust and comprehensive evidence at mesoscopic and millisecond scale for the precise neural deficits of amblyopia in monocular processing and binocular interactions.

### Neural deficits of amblyopia in monocular processing

The locus of amblyopic deficits has been a long-standing question in the literature. In addition to low-level monocular deficits in visual acuity and contrast detection, psychophysical studies also suggest abnormalities in higher visual functions (Levi, 2006, 2020; Sharma et al., 2000), such as second-order detection (Wong et al., 2001), contour and motion integration (Hess et al., 1997; Simmers et al., 2003), shape discrimination (Hess et al., 1999), and attention (Popple & Levi, 2008; Verghese et al., 2019). Neural deficits of amblyopia at multiple levels of the visual hierarchy have also been demonstrated using electrophysiological recordings in animal models (Kiorpes, 2006, 2019), and BOLD fMRI in humans (Anderson & Swettenham, 2006; Lerner et al., 2003; Muckli et al., 2006). The current consensus is that the neural deficit of amblyopia originates in V1 and may lead to additional functional deficits in the downstream visual areas (Anderson & Swettenham, 2006; Kiorpes, 2019; Levi, 2020). However, it remains unclear whether feedforward, feedback, or both processes are disrupted in the hierarchical processing of visual information. Using simple checkerboard stimuli, our 7T fMRI results revealed similar degrees of response deficits in monocular processing from V1 layers receiving thalamic input to the downstream cortical layers and extrastriate visual areas in the ventral stream. These findings support that monocular deficits of amblyopia in the visual cortex are detectable from the outset at the thalamic input layer of V1 and cascade to downstream areas along the feedforward pathway. This is also consistent with the functional and structural abnormalities found in the LGN of the thalamus in human amblyopia (Barnes et al., 2010; Hess et al., 2010; Wen et al., 2021). However, the current findings using simple checkerboard flickers do not preclude the possibility of abnormal feedback processing with more complex visual stimuli.

### The role of suppression in amblyopic deficits and abnormalities in binocular intergration

In addition to the neural deficits in monocular processing, binocular suppression may also play important roles in the visual deficits of amblyopia as suggested by clinical evidence (DeSantis, 2014; Von Noorden, 1996) and psychophysical studies (Baker et al., 2008; Holopigian et al., 1988; Li et al., 2011; Zhou et al., 2013). For example, using dichoptic motion detection, Li and colleagues found a negative correlation between the strength of binocular suppression and the depth of amblyopia (Li et al., 2011). More recent psychophysical evidence showed reduced suppression from the amblyopic eye relative to normal controls (Zhou et al., 2018). In animal models with non-human primate, there was a striking loss of binocular neurons in V1 (Kiorpes, 2019; Kiorpes et al., 1998; Movshon et al., 1987). Stronger than normal binocular suppression of neuronal activity has also been observed in the early visual cortex of strabismic monkeys (Bi et al., 2011), in which the suppression from FE to AE was stronger than from AE to FE (Shooner et al., 2017). However, there has been no clear evidence for abnormal binocular intergration or imbalanced suppression in the very few neuroimaging studies in human amblyopia (Baker et al., 2015; Chadnova et al., 2017; Farivar et al., 2011; Lygo et al., 2021).

In the current study, our 7T fMRI and EEG data provide strong neuroimaging evidence for the role of binocular suppression in the neural deficits of amblyopia and the abnormalities in binocular interactions. In contrast to a flat laminar profile in monocular response deficit, the amblyopic deficit in binocular responses, though underestimated due to partial volume effect, was strongest in the superficial layer of V1 (Fig. 3b), suggesting an additional signal loss due to suppression from the fellow eye by lateral inhibition. SSVEP results from the EEG experiment further demonstrate that the additional response deficit in the binocular condition was a result of imbalanced binocular suppression in the early visual cortex (Fig. 4c/d). This was mainly due to reduced suppression by the amblyopic eye relative to normal controls, consistent with recent psychophysical evidence (Gong et al., 2020; Zhou et al., 2018). In addition, we also found a significant reduction in the intermodulation amplitude. The IM frequency component could only arise from nonlinear combinations of the two eyes’ signals (Baitch & Levi, 1988; Regan & Regan, 1988). Recent studies suggest that the IM component may reflect interocular suppression in binocular vision (Du et al., 2023; Katyal et al., 2018). Therefore, a reduction in the IM amplitude suggests weakened binocular interactions in amblyopic subjects. This finding was also consistent with our finding of reduced suppression from the amblyopic eye (Fig. 4), the loss of binocular neurons found in the monkey studies (Kiorpes, 2019), and a recent study showing a correlation between IM amplitude and the interocular difference in contrast sensitivity (Gu et al., 2020). Based on neuroanatomical and electrophyiological evidence (Cox et al., 2019; Dougherty et al., 2019; Hubel & Wiesel, 1972), binocular neurons mainly exist in the superficial layers of V1, combining the two eyes’ signals through nonlinear normalizations (Mitchell et al., 2023; S.-H. Zhang et al., 2024). Our findings thus filled the gap between psychophysical studies in humans and electrophysiological studies in the animal models about the amblyopic abnormalities in binocular interactions.

### Slower visual processing speed to monocular and binocular stimuli

Delayed visual processing to stimulus presented to the amblyopic eye relative to the fellow eye has been reported in two previous studies (Chadnova et al., 2017; Lygo et al., 2021). However, the difference from normal controls remains unclear. In our current study (Fig. 6), significant phase delays of SSVEPs were observed not only between AE and NE, but also between FE and NE, in both monocular and binocular stimulus conditions. These findings demonstrate slower visual processing speed for both amblyopic and fellow eyes relative to normal controls. The phase delay was unlikely due to reduced neural activity, since the P-pathway biased high SF responses exhibited significant phase delay but much larger response amplitude compared to the M-pathway biased low SF responses. For the control group, binocular stimuli also showed faster response timing than monocular stimuli, suggesting a benefit in visual processing speed with binocular vision. However, the binocular advantage in visual processing speed was not observed in AE or FE responses in amblyopic subjects. This finding provides additional support for amblyopic abnormalities in binocular integration, consistent with the reduction in intermodulation amplitude (Fig. 5). Therefore, our data present robust electrophysiological evidence for delayed visual processing in both the amblyopic and fellow eyes compared to normal controls. Weakened binocular integration, most likely in the superficial layers of V1, further reduced visual processing speed in binocular vision.

## Conclusion

Using a combination of submillimeter 7T fMRI and EEG frequency-tagging methods, we found attenuated and delayed neural activity in monocular processing and binocular interactions in the early visual cortex of amblyopic adults. Our findings support that neural deficits in monocular processing arise from the thalamic input layers of V1, followed by imbalanced binocular suppression and weakened binocular integration in the superficial layers. The precise neural deficits of amblyopia at the mesoscopic and millisecond resolution could help developing more targeted and effective treatments for this common vision disorder.

## Methods and Materials

### Participants

A total of 10 adults diagnosed with unilateral amblyopia (9 anisometropia and 1 strabismus, 26.0 ± 7.90 years of age, 6 females) were enrolled in the fMRI experiment. 37 adult patients diagnosed with unilateral amblyopia (21 anisometropic amblyopia, 4 strabismic amblyopia, 1 mixed amblyopia and 1 deprived amblyopia after cataract surgery, 27.89 ± 5.94 years of age, 25 females) and 15 normal participants (25.20 ± 1.70 years of age, 10 females) were enrolled in the EEG experiment. The research followed the tenets of the Declaration of Helsinki, and all participants gave written informed consent in accordance with procedures and protocols approved by the Human Subjects Review Committee of the Eye and ENT Hospital of Fudan University, Shanghai, China. The participants also gave written informed consent approved by the institutional review board of the Institute of Biophysics, Chinese Academy of Sciences. The best-corrected visual acuity (BCVA) of each subject was assessed by an experienced optometrist with the Snellen chart. Cover and alternative cover testing, duction and version testing, intraocular pressure testing, slit-lamp testing, indirect fundus examination after pupil dilation, and optometry were performed for each subject. The BCVAs of all amblyopic eyes was > 0.1 logMAR and < 1.0 logMAR, and the BCVA of all fellow eyes was ≤ 0.1 logMAR or better. Amblyopia patients were required to be free from a history of intraocular surgery, any eye diseases, and any systemic diseases known to affect visual function (e.g., migraine, congenital color deficiencies). None of amblyopia subjects underwent amblyopia treatment including patching treatment. All strabismic amblyopia patients had undergone strabismus surgery at least one year previously and had normal eye position and steady fixation. Eye dominance in normal subjects was determined by instructing the subjects to look at a distant letter with both eyes open through a hole between their hands, and then with their eyes alternatively closed. The eye that could still see the target was the dominant eye (hole-in-card method). The clinical details of all patients were listed in supplementary table S1.

### Stimuli and procedures

In the fMRI experiment, a full contrast checkerboard (30-by-22.5 degrees of visual angle, 0.26 degrees checker size) counterphase flickering at 8 Hz was presented through a pair of MR-compatible goggles (NordicNeuroLab). The checkerboard flickers were presented to the amblyopic eye, the fellow eye, or both in separate blocks (18 s), interleaved with 12-s fixation blocks. Each run lasted 282 s, comprising 9 stimulus blocks (3 for each condition) and 10 fixation blocks. The order of stimulus conditions was counterbalanced across the 8 runs. During the experiment, subjects were instructed to keep fixating a central point. Both eyes wore glasses to correct for refractive errors.

In the EEG experiment, visual stimuli were low- and high-pass filtered natural scene images equalized in brightness and root mean square contrast. Stimuli were on and off flickered at two temporal frequencies, 7.2 and 8 Hz, delivered one to each eye during monocular and binocular stimulus presentations. The association of tagging frequencies and eyes was balanced within each subject: 7.2 Hz for the left eye and 8 Hz for the right eye, and vice vera. Therefore, the EEG stimulation included a total of 2 SFs * 3 ocularities * 2 TFs = 12 stimulus conditions. Stimuli were dichoptically presented with a pair of base-out prism glasses and a cardboard divider splitting the left and right visual fields. A pair of circular frames with mosaic patterns outside the frames were used to assist fusion between the two eyes. Participants pressed a button to start a trial. They were instructed to keep fixation and avoid blinking as much as possible during the 6-second period of stimulus presentation. There were 20 stimulus blocks in the experiment, each had 12 stimulus conditions in a pseudo randomized order.

### MRI data acquisition

MRI data were acquired with a 7T scanner (Siemens Magnetom) using a custom 32-channel receive 4-channel transmit open-face head coil (Fig. 1b; the “Visual coil”) in the Beijing MRI Center for Brain Research (BMCBR). A bite bar was used to reduce head motion. The open-face design of the head coil enabled using both goggles and the bite bar. The gradient coil has a maximum amplitude of 70mT/m, 200 μs minimum gradient rise time, and 200 T/m/s maximum slew rate.

High resolution functional data were collected using a T2*-weighted 2D gradient-echo EPI sequence (0.8 mm isotropic voxels, TR = 2000 ms, TE = 23 ms, nominal flip angle = 78°, field of view 128 × 128 mm, 31 oblique-coronal slices covering the posterior part of the brain, receiver bandwidth 1157 Hz/pix, 6/8 phase partial Fourier, GRAPPA acceleration factor = 3, phase encoding direction from A to P). Two dummy scans were collected at the beginning of each run. Five EPI images with reversed phase encoding direction (P to A) were also acquired for EPI distortion correction before each run. T1-weighted anatomical volumes were acquired using a MP2RAGE sequence (0.7 mm isotropic voxels, FOV = 224×224 mm, 256 sagittal slices, TE = 3.05 ms, TR = 4000 ms, TI1 = 750 ms, flip angle = 4°, TI2 = 2500 ms, flip angle = 5°, bandwidth = 240 Hz/pix, partial Fourier = 7/8, GRAPPA = 3). For cortical segmentations, we collected whole-brain T1w-MP2RAGE images in a separate session using a 32-channel receive 1-channel transmit head coil (the “NOVA coil“; Nova Medical).

### MRI data preprocessing

MRI data were preprocessed using AFNI (Cox, 1996) and the mripy package developed in our lab (https://github.com/herrlich10/mripy). EPI volumes were corrected for slice timing, susceptibility distortion (blip-up/down method), head motion (6 parameters rigid body), and rescaled to percent signal change. To minimize the loss of spatial resolution, all spatial transformations were combined and applied in a single interpolation step (sinc method). Slow baseline drift and motion parameters were included as regressors of no interest in addition to the stimulus regressors in a general linear model (GLM). A canonical HRF (BLOCK4 in AFNI) was used in the GLM analysis.

### Laminar analysis

The outline of the data analysis pipeline was shown in Fig. S1. The whole-brain anatomical volume was segmented into white matter, gray matter and CSF, based on which pial and white matter surfaces were reconstructed using FreeSurfer (v6.0) (Fischl, 2012) with -hires option. Results were visually inspected and manually edited. To prevent the surface-to-volume projection from creating “holes” in high resolution volumes, high-density meshes were generated by 4-fold up-sampling from the default ones. The anatomical volume from the NOVA coil as well as the reconstructed surfaces were first aligned to the T1 image from the Visual coil, and then to the mean EPI image. Relative cortical depths (zero at pial and one at white matter surface) of gray matter voxels were estimated using a equivolume method (Waehnert et al., 2014). Surface curvatures were considered in the depth estimation by taking the ratio of neighborhood areas for a pair of corresponding vertices on the pial and white matter surfaces. A set of intermediate surfaces were generated at various equivolume distances and the depth value for each voxel was estimated by linear interpolation between two nearest surfaces (mripy_align_anat.ipy from mripy). The superficial, middle and deep layers were defined to each occupy 1/3 of the gray matter volume (Balaram et al., 2014; de Sousa et al., 2010; Liu et al., 2021). Cortical depth-dependent responses were averaged across all voxels in a layer compartment.

To reduce the influence from large veins on the BOLD laminar profiles, voxels contaminated by large veins were excluded in a column-wise manner. The bias-field corrected mean EPI intensity and averaged BOLD responses were mapped onto the cortical surface. Vertices with an EPI intensity lower than 70% of the averaged intensity or with a BOLD signal change over 10% were identified as large veins. These vertices were subsequently mapped back to the volumetric space to form a mask that encompassed all voxels within the same column. Voxels within this mask were excluded from subsequent analysis.

### ROI definition

Anatomical ROIs for early (V1, V2, V3) and intermediate (V4, LO, V3ab) visual areas were defined using a retinotopic atlas based on 7T fMRI data from the Human Connectome Project (Benson et al., 2018). In the monocular condition, vertices exhibiting significant visual activation (mean across AE, FE and binocular, p < 0.05 uncorrected) within the anatomical atlas were included in the analyses. In the binocular condition, ocular biased vertices in V1 were defined by the response difference between AE and FE in the monocular condition (10% vertices from each side of the distribution). In a supplementary analysis, we used a split-half-run approach to define the same ROIs for analyzing monocular and binocular responses (Fig. S2). Ocular biased vertices (sorted by AE-FE beta) with significant visual activations (mean response p < 0.05 uncorrected) in odd runs were defined as the surface ROIs for analyzing monocular and binocular responses in even runs. We then used the data from even runs as an independent localizer to extract the responses from odd runs. Results from the two split-half-run analyses were averaged.

### EEG data collection

EEG data were collected with a 64-channel system (BrainVision actiCHamp), digitized at 1000 Hz. The ground electrode was located in front of AFz, and the reference electrode between Fz and Cz. Electrode impedance was maintained below 8 kΩ throughout the experiment. A DELL S2721DGF monitor was used for stimulus presentation, setup at 144 Hz refresh rate and 2560*1440 resolution. The distance between the monitor and the eyes of participants was 85 cm.

### EEG data analysis

EEG data were preprocessed using EEGLAB in Matlab. For SSVEP amplitude analysis, a bandpass filter between 1 and 30 Hz was applied on the raw EEG time series. To avoid the transient response from stimulus onset, we used the last 5 seconds of data in each trial for further analysis. Mean time series were generated by averaging across all trials in each condition. A spatial Laplacian filter was used to increase the signal-to-noise ratio, with O1, Oz, O2 and PO3, POz, PO4 as the central electrodes, and P5, P3, P1, Pz, P2, P4, P6 and PO7, PO8 as the surround electrodes. The differential signal between the means of central and surround electrodes was fast Fourier transformed to obtain the amplitude spectrum. Amplitudes of the first and the second order harmonics were summed together. The amplitudes of SSVEPs to the 7.2 and 8 Hz stimuli were averaged for each condition. For the phase analysis, a bandpass filter with a 6-9 Hz frequency range was applied to the raw time series. EEG data from 1251 to 5000 ms (integral multiples of periods at both stimulus frequencies) after stimulus onset were used. A least squares filtering method was used to calculate the phase of mean time series at the stimulus frequency.

### Statistics

Statistical tests were performed using Pingouin (v0.5.4) and JASP (v0.14.1). If not otherwise specified, the comparison of mean responses between conditions, layers, or groups were tested for significance using ANOVA followed by two-tailed t-test. If multiple measurements were taken from the same group under different conditions or from different ROIs, repeated measures ANOVA and paired t-test were used. The follow-up tests were corrected for multiple comparisons using the Holm method, unless there were only three levels followed by a significant one-way ANOVA, or two levels followed by a significant 2×2 interaction, in which cases a correction is not necessary (Cohen, 2008; Levin et al., 1994). Partial eta squared and Cohen’s d were reported as the effect size for ANOVA and t-test, respectively. When comparing BSI within and between groups, non-parametric methods (Wilcoxon signed-rank test for paired tests and Wilcoxon rank-sum test for independent samples test) were used due to the large difference in variance. Rank-biserial correlation (RBC) was reported as effect size. Error bars in the bar-plots indicate 95% confidence interval, which was generated using a bootstrap method (seaborn v0.12.2).

We adopted a multivariate approach in detecting data outliers. For each set of comparisons, the measurements from the same subject were considered as a single point in a multidimensional space, and the points whose likelihood was less than 1% (assuming an underlying multivariate normal distribution) were flagged as outliers (encircled by red outlines in Fig. 4c/d and Fig. 5) and excluded from subsequent statistical tests. The likelihood was obtained by evaluating the Mahalanobis distance (squared) of the point against a chi-squared distribution with degrees of freedom equal to the number of dimensions. The phase differences in Fig. 6 were tested using the Circular Statistics Toolbox (v1.21) in Matlab (Berens, 2009). ‘circ_mtest’ was used to test the within-subject phase delay between stimulus conditions. ‘circ_wwtest’ was used to test the phase delay between AE/FE and NE.

## Data and code availability

Data and code to reproduce the main findings of this study will be made available upon publication.

## Acknowledgement

This study was supported by Ministry of Science and Technology of China (https://en.most.gov.cn/) STI2030-Major Projects (2022ZD0211900 to P.Z., 2021ZD0204200 to C.Q.), National Natural Science Foundation of China (https://www.nsfc.gov.cn/english/site_1/index.html, 31930053 to P.Z., 32000787 to C.Q.),

Project of State Key Laboratory of Ophthalmology, Optometry and Vision Science, Wenzhou Medical University (http://skloovs.ac.cn/, J02-20210203 to P.Z.), Youth Innovation Promotion Association CAS (https://english.cas.cn/, 2021089 to C.Q.). Shanghai Science and Technology Innovation Action Plan, (https://svc.stcsm.sh.gov.cn/, 22ZR1410200 to W.W.).

## Supplementary information

**Table S1.** Clinical details of amblyopic subjects.

| Subject ID | Age | Gender | Handedness | BCVA<br>(logMAR) | Type of amblyopia | Eye alignment |
| --- | --- | --- | --- | --- | --- | --- |
| 7T P01 | 37 | F | right | 1.0 | anisometropia | orthotropia |
| 7T P02 | 28 | M | right | 1.0 | anisometropia | orthotropia |
| 7T P03 | 19 | M | right | 1.0 | anisometropia | orthotropia |
| 7T P04 | 20 | F | right | 0.7 | anisometropia | orthotropia |
| 7T P05 | 23 | M | right | 0.5 | strabismic | Orthotropia (post surgery) |
| 7T P06 | 16 | M | right | 1.1 | anisometropia | orthotropia |
| 7T P07 | 33 | F | right | 0.5 | anisometropia | orthotropia |
| 7T P08 | 17 | F | right | 0.4 | anisometropia | orthotropia |
| 7T P09 | 33 | F | right | 0.3 | anisometropia | orthotropia |
| 7T P10 | 34 | F | right | 0.8 | anisometropia | orthotropia |
| EEG S01 | 29 | F | Left | 0.4 | anisometropia | orthotropia |
| EEG P02 | 25 | M | Left | 0.3 | anisometropia | orthotropia |
| EEG P03 | 26 | F | Right | 0.6 | anisometropia | orthotropia |
| EEG P04 | 34 | F | Right | 0.7 | anisometropia | orthotropia |
| EEG P05 | 23 | M | Left | 0.7 | anisometropia | orthotropia |
| EEG P06 | 21 | F | Left | 0.3 | anisometropia | orthotropia |
| EEG P07 | 25 | F | Left | 0.4 | anisometropia | orthotropia |
| EEG P08 | 30 | F | Right | 0.4 | anisometropia | orthotropia |
| EEG P09 | 30 | F | Right | 0.3 | strabismus | orthotropia (post surgery) |
| EEG P10 | 25 | F | Left | 0.5 | anisometropia | orthotropia |
| EEG P11 | 28 | M | Left | 0.3 | anisometropia | orthotropia |
| EEG P12 | 30 | F | Right | 0.4 | anisometropia | orthotropia |
| EEG P13 | 30 | F | Right | 0.5 | strabismus | orthotropia (optical correction) |
| EEG P14 | 26 | M | Right | 0.8 | anisometropia | orthotropia |
| EEG P15 | 18 | F | Right | 0.2 | anisometropia | orthotropia |
| EEG P16 | 29 | M | Left | 0.1 | anisometropia | orthotropia |
| EEG P17 | 26 | F | Left | 0.5 | anisometropia | orthotropia |
| EEG P18 | 34 | F | Left | 0.3 | anisometropia | orthotropia |
| EEG P19 | 25 | F | Left | 0.5 | anisometropia | orthotropia |
| EEG P20 | 37 | F | Right | 0.7 | anisometropia | orthotropia |
| EEG P21 | 29 | F | Left | 0.5 | anisometropia | orthotropia |
| EEG P22 | 21 | M | Left | 0.7 | anisometropia | orthotropia |
| EEG P23 | 18 | F | Left | 0.7 | mixed | orthotropia (post surgery) |
| EEG P24 | 23 | M | Left | 0.7 | deprived | orthotropia (post surgery) |
| EEG P25 | 34 | F | Left | 0.7 | anisometropia | orthotropia |
| EEG P26 | 21 | F | Left | 0.4 | anisometropia | orthotropia |
| EEG P27 | 37 | M | Right | 0.6 | anisometropia | orthotropia |
| EEG P28 | 27 | F | Right | 0.5 | anisometropia | orthotropia |
| EEG P29 | 24 | F | Right | 0.3 | anisometropia | orthotropia |
| EEG P30 | 23 | M | Right | 0.8 | anisometropia | orthotropia |
| EEG P31 | 22 | M | Right | 0.5 | anisometropia | orthotropia |
| EEG P32 | 22 | F | Left | 0.4 | anisometropia | orthotropia |
| EEG P33 | 35 | M | Left | 0.4 | strabismus | orthotropia (optical correction) |
| EEG P34 | 30 | F | Left | 0.7 | anisometropia | orthotropia |
| EEG P35 | 33 | F | Right | 0.5 | anisometropia | orthotropia |
| EEG P36 | 39 | M | Right | 0.2 | strabismus | orthotropia (optical correction) |
| EEG P37 | 43 | F | Left | 0.5 | anisometropia | orthotropia |

**Figure S1.**
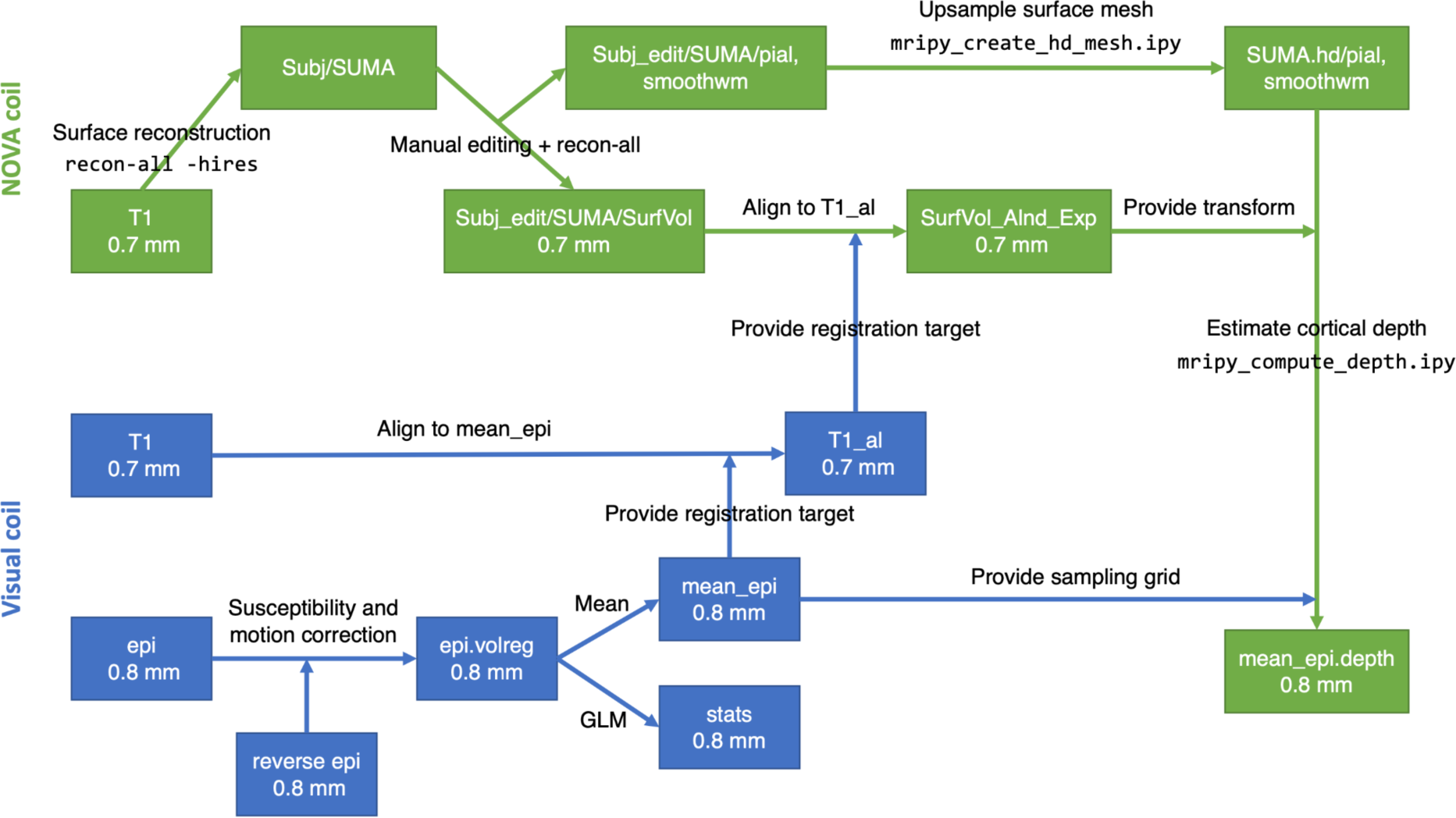
Overview of the anatomical and functional MRI data processing pipeline. Blue and green boxes indicate datasets derived from the data acquired using the Visual and NOVA coils, respectively.

**Figure S2.**
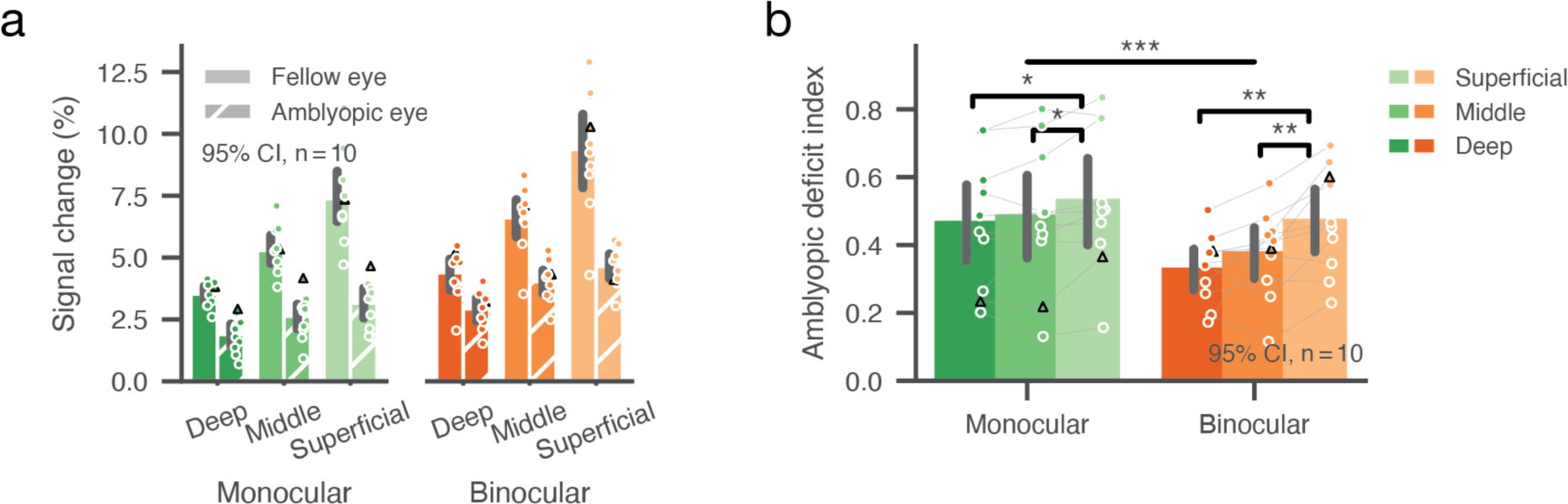
Cortical depth-dependent fMRI response (a) and amblyopic deficit indices (b) in V1 for monocular and binocular conditions using a split-half approach. To compare V1 responses in monocular and binocular conditions within the same set of voxels selected using independent data, we split fMRI data into two halves (odd runs and even runs). We selected AE- and FE-biased vertices using the first half, and then evaluated the mean response with the surface ROI using the second half of the data. The superficial layers now also showed a slightly larger amblyopic deficit than the deeper layers during monocular stimulation, but the difference was more prominent during binocular stimulation as suggested by the highly significant layer-by-condition interaction. Conventions were identical as in Fig. 3a/b.

**Figure S3.**
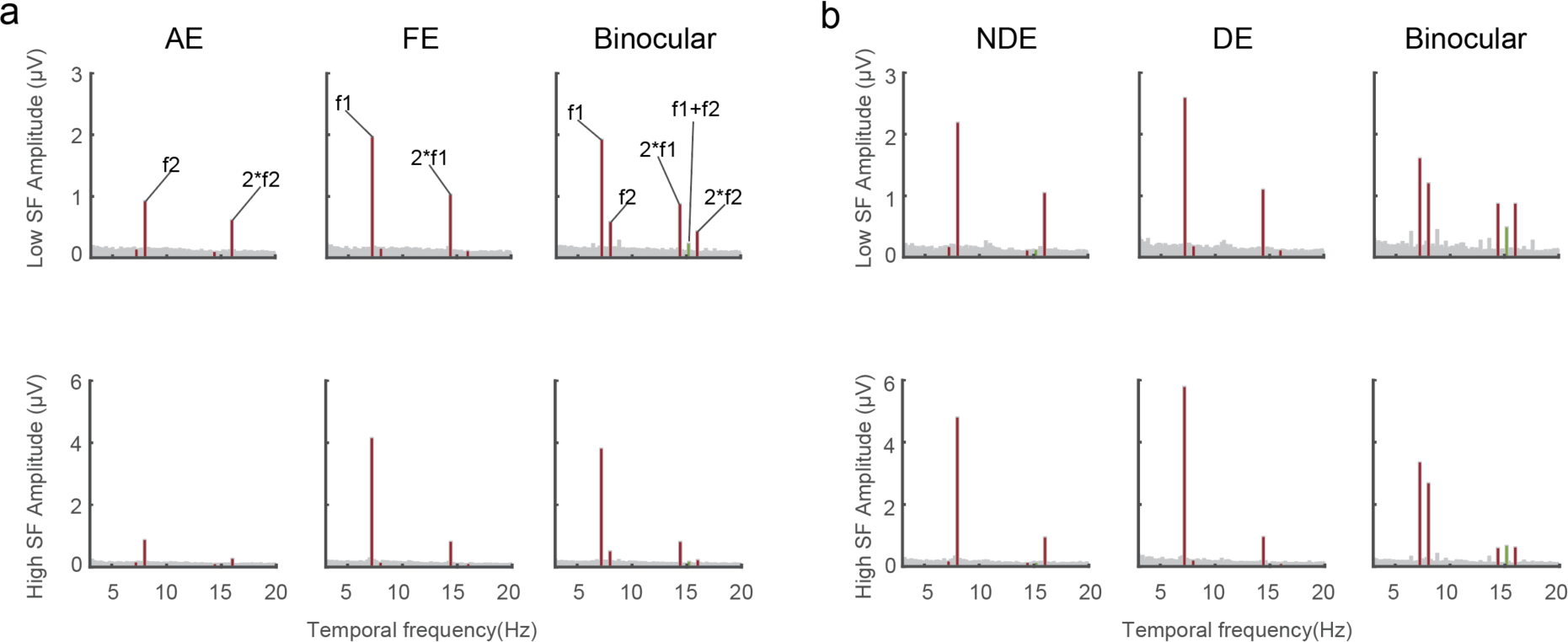
**(a)** Group-averaged amplitude spectrums to stimuli presented to AE (f2) and FE (f1) in amblyopic subjects. **(b)** Group-averaged amplitude spectrums to stimuli presented to NDE (f2) and DE (f1) in normal controls.

